# Large scale changes in host methylation patterns induced by IncA/C plasmid transformation in *Vibrio cholerae*

**DOI:** 10.1101/399964

**Authors:** Ruibai Wang, Kanglin Wan

## Abstract

DNA methylation is a central epigenetic modification and has diverse biological functions in eukaryotic and prokaryotic organisms alike. The IncA/C plasmid genomes are approximately 150kb in length and harbour three methylase genes, two of which demonstrate cytosine specificity. Transformation of the *Vibrio cholerae* strain C6706 with the IncA/C plasmid pVC211 resulted in a significant relabelling of the methylation patterns on the host chromosomes. The new methylation patterns induced by transformation with IncA/C plasmid were accepted by the restriction enzymes of the host’s restriction modification (RM) system. These data uncover a novel mechanism by which plasmids can be compatible with a host’s RM system and suggest a possible reason that plasmids of the IncA/C family are broad-host-range.

**Author summary:** Antibiotic resistance of bacteria is a growing serious problem worldwidely and the horizontal transfer of multi-drug resistance genes mediated by plasmids within and between species of bacteria is the main reason. In the researches of multi-drug resistance of *Vibrio cholerae*, I have isolated several IncA/C plasmids. What impressed me most is their ability to accumulate the resistant genes. Moreover, they can transfer with high frequency and are stable in several bacterial species. There are at least three Tra regions on the IncA/C plasmid which containing components of the Type 4 Secretion System and are important for conjugative transfer of plasmids. So the horizontal transfer ability of IncA/C plasmids is reasonable. There are three methylase genes on the small genome of IncA/C plasmids, which demonstrate cytosine specificity and are seldom in bacteria. Their modification target and roles are interesting. Here, we analysed the methylation profiles of the host *V. choerae* induced by the plasmid pVC211 and found that they were completely changed. In addition to replicons, this may be a novel mechanism that plasmid cross the barrier of the host’s RM system and become broad-host range. Changing the activity of methylase in IncA/C plasmids may be a new way to affect the stability of IncA/C plasmids to eliminate these multidrug-resistant plasmids from bacteria.

## 1 Introduction

IncA/C family plasmids are important carriers of multi-drug resistance genes in plasmid-incompatible groups (Inc) and mediate the dissemination of multi-drug resistance in bacteria [1,2]. Both of the plasmid pIP1202, which has high resistance to at least eight antibiotics isolated from *Yersinia pestis* IP275 in 1995 [3], and the plasmid of NDM-1 super-resistant bacteria in India, Bangladesh, Pakistan, Britain and the United States in August 2010 [4] are IncA/C family plasmids and have aroused great concern in public health and bioterrorism. IncA/C plasmids have great ability to accumulate antibiotic resistant genes. There are many resistance genes of rifampicin, erythromycin, streptomycin, chloramphenicol, sulfonamides and disinfectants routinely harbored on the plasmids [5], as well as a variety of beta-lactamase genes.

For example, the pNDM-1_Dok01, pNDM102337, pNDM10469, pNDM10505, pNDMCFuy, pNDM-KN and pNDM-US plasmids of *Escherichia coli* and *Klebsiella pneumoniae*. The pVC211 plasmid we isolated in *Vibrio cholerae* has 16 antibiotic resistance related genes, including five macrolide resistant genes involving in three mechanisms [6]. The IncA/C plasmid made 99% of *Vibrio cholerae* O139 strains resistant to more than three antibiotics in which 47% were resistant to eight antibiotics [7].

IncA/C plasmids are a kind of plasmid with wide host range. They exist and transfer horizontally in many bacterial genera such as *E. coli*, *Salmonella, Enterobacter*, *Klebsiella, V. cholerae*, *Yersinia*, *Pantoea, Edwardsiella*, *Citrobacter freundii*, *Photobacteriumdamselae, Aeromonas*, *Xenorhabdus nematophila, Providencia,* with the transformation efficiency as high as 10^-1^ to 10^-2^. The horizontal transfer of multi-drug resistance genes mediated by plasmids within and between species of bacteria are the main reason for the growing problem of drug resistance worldwide. In addition, the skeleton of IncA/C plasmid also widely distribute in multidrug-resistant agricultural pathogens. Strains carrying IncA/C plasmids have been isolated from beef, chicken, turkey and pork. Reports showed that the *bla*CMY-2 gene mediated by IncA/C plasmid and conferring resistant to cephalosporin could be identified in human *E. coli* and *Klebsiella pneumoniae* isolates several years after the prevalence of the gene in edible animals [8]. Moreover, 1% of *E. coli* strains isolated from healthy people who had never taken antibiotics were positive for *repA* / *C* gene, the replicator of IncA / C plasmid [9,10]. This mobile reservoir of drug resistance determinants can transmit multi-drug resistance phenotypes from foodborne pathogens to human pathogens, conducting the using impact of veterinary drug to human drug resistance profiles, and has special public health implications.

IncA/C plasmids are large, conjugative plasmids about 150kb in length. On the plasmid backbones, there are three methylation-related genes: *dcm1*, *dcm2*, and *dcm3.* These genes are 1626 bp, 1428 bp and 924 bp in length and are identical to NCBI reference sequences WP_000201432.1, WP_000936896.1 and WP_000501488.1, respectively. The *dcm1* gene encodes Gammaproteobacteria cytosine-C5 specific DNA methylases (MTase) of the Pfams PF00145 family. The *dcm2* gene is a DNA cytosine methyltransferase with an AdoMet_MTases (cl17173) conserved domain. The *dcm3* gene is a DNA modification MTase that lack any known conserved domain. These three methylation genes are broadly conserved in other IncA/C_2_ plasmids, although a few of plasmids have lost the *dcm2* genes due to the deletion event associated with the ARI-B resistance island [2]. There are six methylation-related genes in the host genome of *V. cholerae*; three adenine-specific MTase genes (*dam*: VC1769, VC2118, and VC2626), two rRNA MTases (VC2697 and VCA0627), and an orphan cytosine MTase, vchM (VCA0158). Generally, DNA (cytosine-5-) MTases (Dcm) recognized a CCWGG motif and covalently add a methyl group at the C5 position of the second cytosine C, while Dam introduce a methyl group at the N6 position of the adenine in GATC motifs. The MTases of *V. cholerae* chromosomes are mainly Dam MTases and the only Dcm MTase, VchM, recognizes a novel target, the first 5’C on both strands of the palindromic sequence 5’-RCCGGY-3’, and leave all 5’-CCWGG-3’ sites unmethylated in *V. cholerae* [11]. These data suggest that changes in methylation mediated by IncA/C plasmids may be quite different from that of the host chromosomes.

DNA methylation is a central epigenetic modification in various cellular processes including DNA replication and repair [12], transcriptional modulation, decreasing transformation frequency, the stability of short direct repeats in certain bacteria, and is necessary for site-directed mutagenesis [13]. DNA methylation functions either alone or as part of the bacterial restriction modification (RM) system, protecting the host from infection by foreign DNA and bacteriophage by degrading non-methylated DNA with sequence-specific restriction enzymes. Though the role of IncA/C plasmids in drug resistance is well elucidated, the role of the *dcm* genes on A/C_2_ plasmids remains largely unexplored. In this study, the IncA/C plasmid pVC211 (148,456bp) [6], was conjugated into *V. cholerae* strain C6706. Sequencing of genomic DNA provided a view of the change in the methylation patterns of the host that were induced by this plasmid.

Common methylation modifications cells are 6-methyladenine (^m6^A), 4-methylcytosine (^m4^C), and 5-methylcytosine (^m5^C) [14]. There are two ways to measure DNA methylation on a genome-wide scale: bisulfite sequencing and single-molecule, real-time (SMRT) sequencing. Bisulfite sequencing is commonly used to detect ^m5^C. In this method, treatment of DNA with bisulfite converts cytosine residues to uracil, leaving 5-methylcytosine residues unaffected. The methylation status of a fraction of DNA can be determined by the analysis performed on such alterations. SMRT sequencing identifies bases using kinetic information recorded during each nucleotide addition step, which can be used to distinguish modified and native bases by comparing with the kinetic reference for dynamics without modification. Different epigenetic modification types have unique kinetic signatures and can be inferred, including the methylation at each GATC site. But because the ^m5^C modification produces weaker and somewhat more diffuse SMRT signals and requires 10 fold sequencing coverage, only all predicted ^m6^A and ^m4^C MTases are able to be detected unambiguously in this way [14,15]. In this study, the two complementary methods were used to obtain a global view of methylation change induced by IncA/C plasmids in *V. cholerae*.

## 2 Results

### 2.1 Methylomes determined by SMRT sequencing

The general genome information and methylation analysis by the P_modification detection model in the SMRT sequencing are listed in the Table 1. The two samples, C6706 and CV2, produced 998,990,159 bp and 1,097,369,564 bp polymerase reads respectively, and the averages of the sequencing coverage were ∼250×. In the C6706 genome, 1,476,420 methylation sites were detected and 1,490,035 methylation sites were detected in the CV2. The total methylation modification ratios of both samples were similar, 36.6% and 36.94%. The highest proportion of the methylation types in both C6706 and CV2 were bases reported as ‘modified bases’, which were suspected sites failing to show any definitive methylation patterns by SMRT analysis.

**Table 1.** The general genome and methylation information derived from SMRT sequencing.

| Sample | Polymerase reads data (Mb) | Subreads data (Mb) | Genome size (bp) | Genome map rate (%) | Methyl type | CDS number | Intergenic number | Total number | Modifications ratio (%) |
| --- | --- | --- | --- | --- | --- | --- | --- | --- | --- |
| C6706 | 998 | 898 | 4033464 | 95.43 | m<sup>4</sup>C | 151109 | 4362 | 155471 | 3.85 |
|  |  |  |  |  | m<sup>6</sup>A | 41007 | 1232 | 42239 | 1.05 |
|  |  |  |  |  | untyped |  |  | 1277848 | 31.68 |
| Total |  |  |  |  |  |  |  | 1476420 | 36.60 |
| CV2 | 1097 | 997 | 4033464 | 89.48 | m<sup>4</sup>C | 146542 | 4209 | 151472 | 3.76 |
|  |  |  |  |  | m<sup>6</sup>A | 40686 | 1198 | 41952 | 1.04 |
|  |  |  |  |  | untyped |  |  | 1296611 | 32.14 |
| Total |  |  |  |  |  |  |  | 1490035 | 36.94 |

There were 4013 annotated genes on the *V. cholerae* chromosome, including 2775 functional genes, 25 rRNA genes, and 94 tRNA genes distributed on chromosome I and 1115 functional genes and 4 tRNA genes distributed on chromosome II (Table 2). The methylation of the two chromosomes of *V. cholerae* was considerably different. The genes on chromosome I were generally methylated, dominated by ^m4^C-^m6^A double methylation, and sites involving ^m4^C and ^m6^A accounted for 84.7% (3401/4013) of the total genes. On the other hand, more than half of the genes on chromosome II exhibited non-^m4^C non-^m6^A methylation. In particular, the methylation of the tRNA gene was the most significantly different between the two chromosomes, with 92 of the 94 genes on chromosome I involved in the methylation of ^m4^C or ^m6^A, while on chromosome II, only one of the four genes, VC_At2, was ^m4^C methylated. Chromosome II was a mega-plasmid ameliorated to the host chromosome as a result of its long-standing presence in the host lineage [18], and differences in the methylation pattern of chromosome II still demonstrated its heterogeneity with chromosome I. However, there was no difference between C6706 and CV2 in terms of methylation differences between the two chromosomes.

**Table 2.** The methylation patterns of the two chromosomes of C6706 and CV2.

| Sample | Chromosome | Gene code | Total | none<br>m<sup>4</sup>C/m<sup>6</sup>A | m<sup>4</sup>C | m<sup>6</sup>A | m<sup>4</sup>C and<br>m<sup>6</sup>A |
| --- | --- | --- | --- | --- | --- | --- | --- |
| C6706 | I | VC | 2775 | 14 | 157 | 2 | 2602 |
|  |  | VC_r | 25 | 0 | 8 | 0 | 17 |
|  |  | VC_t | 94 | 2 | 48 | 0 | 42 |
|  | II | VC_A | 1115 | 591 | 177 | 8 | 339 |
|  |  | VC_At | 4 | 3 | 1 | 0 | 0 |
|  | Total |  | 4013 | 610 | 391 | 10 | 3000 |
| CV2 | I | VC | 2775 | 16 | 165 | 6 | 2588 |
|  |  | VC_r | 25 | 0 | 6 | 0 | 19 |
|  |  | VC_t | 94 | 2 | 52 | 1 | 39 |
|  | II | VC_A | 1115 | 590 | 186 | 8 | 331 |
|  |  | VC_At | 4 | 3 | 1 | 0 | 0 |
|  | Total |  | 4013 | 611 | 410 | 15 | 2977 |

The he methylation sites on the genomes of C6706 and CV2 showed significant changes of methylation after conjugation of the IncA/C plasmid (Table 3). A total of 124,571 sites, 85,972 ^m4^C and 38,599 ^m6^A, in 3341 genes kept the same methylation type between the two samples. C6706 had about 67,545 methylation sites that were unique from CV2, and CV2 had about 62,657 unique methylation sites. These differential sites accounted for 37.46% and 35.16% of the C6706 and CV2 methylation sites respectively, involving 81% of the genes (97% of chromosome I and 40% chromosome II). Such a large-scale methylation change made it impossible to discern any modification preference of gene function. Among the genes with differential methylation sites, the methylation modes of 116 genes changed significantly (Supplementary Table 1); for exemple, VC_0025 changed from ^m4^C-^m6^A double methylation to ^m4^C single methylation, VC_A0698 changed from ^m4^C methylation to ^m6^A methylation, and VC_1030 changed from ^m4^C methylation to non-^m4^C non-^m6^A methylation. Though 64 genes were hypothetical proteins with unknown function, other differentially methylated genes involved many key processes in the lifecycle of the *V. cholerae* host. These genes included 1 LuxR family, 3 LysR family transcriptional regulators, 5 regulatory proteins related to phosphoglycerate transport, cold shock, sigma-54 dependent transcription, response and mannitol metabolism, Hcp-1 involved in chromosome segregation, 7 transport and binding proteins related to Na+/H+, vibriobactin and enterobactin transportation and 5 proteins related to protein synthesis and cell fate.

**Table 3.** The common and differential methylation sites between C6706 and CV2.

|  | Sample | Chromosome | Gene code | methyl site number | m<sup>4</sup>C | m<sup>6</sup>A | Gene number |
| --- | --- | --- | --- | --- | --- | --- | --- |
| Common methyl site |  | I | VC | 117587 | 80978 | 36609 | 2744 |
|  |  |  | VC_r | 2116 | 1697 | 419 | 25 |
|  |  |  | VC_t | 384 | 314 | 70 | 85 |
|  |  | II | VC_A | 4481 | 2980 | 1501 | 486 |
|  |  |  | VC_At | 3 | 3 | 0 | 1 |
|  |  | total |  | 124571 | 85972 | 38599 | 3341 |
| Differential methyl sites | C6706 | I | VC | 63533 | 61330 | 2203 | 2715 |
|  |  |  | VC_r | 1149 | 1093 | 56 | 23 |
|  |  |  | VC_t | 248 | 236 | 12 | 79 |
|  |  | II | VC_A | 2613 | 2476 | 137 | 464 |
|  |  |  | VC_At | 2 | 2 | 0 | 1 |
|  |  | total |  | 67545 | 65137 | 2408 | 3282 |
|  | CV2 | I | VC | 58801 | 56919 | 1882 | 2700 |
|  |  |  | VC_r | 1176 | 1082 | 94 | 24 |
|  |  |  | VC_t | 237 | 231 | 6 | 82 |
|  |  | II | VC_A | 2440 | 2335 | 105 | 447 |
|  |  |  | VC_At | 3 | 3 | 0 | 1 |
|  |  | total |  | 62657 | 60570 | 2087 | 3254 |

Analysis of motif sequence using P_motifFind model of SMRT further revealed the effects of plasmid MTase on methylation status of the host chromosomes (Table 4). Eleven and ten motifs were detected in C6706 and CV2 respectively, but only 6 motifs were commonly detected in both samples and the numbers of these motifs differed significantly between the two samples. Four motifs found in C6706, AGKNNNNW, GADNDGCG, GARVNNDG, TVVVNNDG, were changed to ABNBMVBW, GANNDBBG, GARVNRNG and TNVVNNDG in CV2 respectively. The DCAGVHRNG motif present in C6706 disappeared in CV2. These data indicated that plasmid transformation not only changed the methylation types of the host chromosomes, but also fundamentally changed the sequence characteristics for the identification sites of the MTase.

**Table 4.**
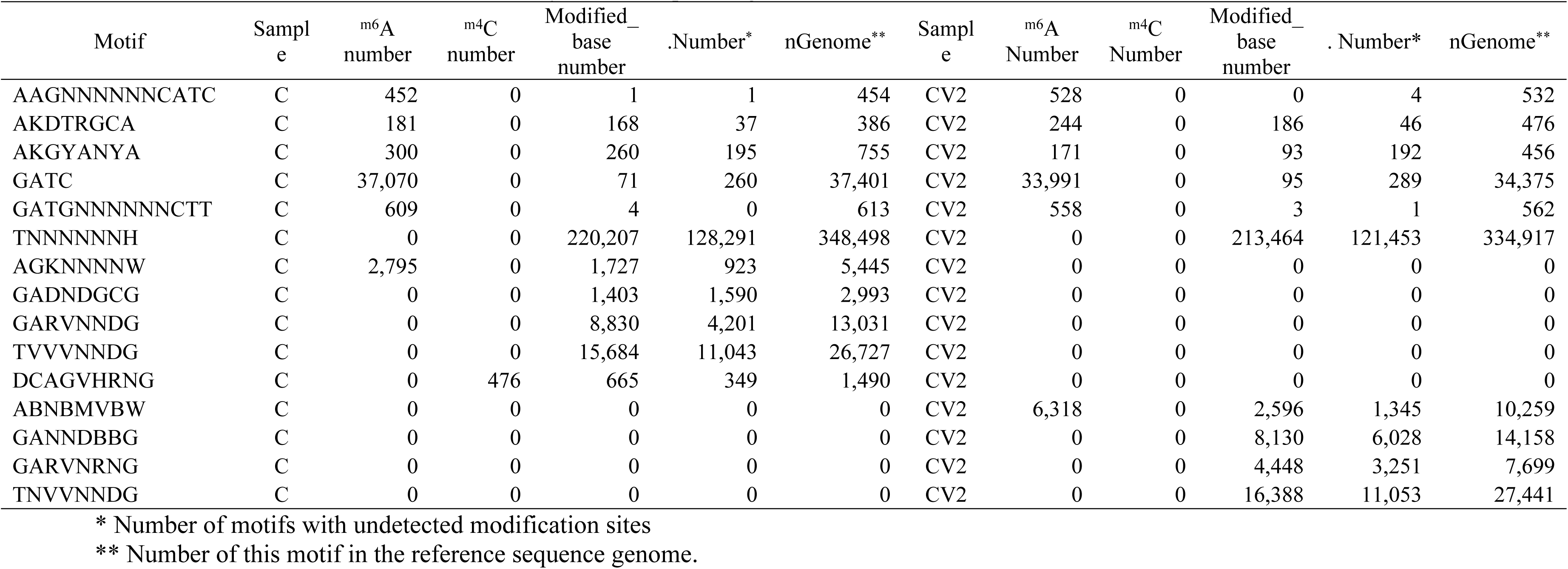
The motifs in C6706 and CV2 determined by SMRT sequencing.

### 2.2 ^m5^C methylation in bisulfite sequencing

Sample C6706 and CV2 produced 16,196,082 bp and 16,232,236 bp clean reads respectively. The mapping rates were 96.01% and 91.78%, the bisulfite conversion rates were 99.65% and 99.62%, and the average depths were 362.12× and 345.69× respectively. We detected 6,965 methylcytosines in C6706 and 7,321 methylcytosines in CV2 from a total of 1,915,376 cytosines in the *V. cholerae* genome. The average methylation levels of different types of C bases (mCG, mCHG and mCHH, where H = A, C or T) increased in CV2 comparing to C6706, and on a whole genome scale were 0.33%, 1.74% and 0.43% in C6706 and 0.34%, 1.81% and 0.44% in CV2. Moreover, the graphs of the methylation levels also showed a slightly increasing trend from C6706 to CV2 (Fig. 1A). For example, 50% of mCHH sites were 10% methylated in C6706; this was increased in CV2 as 59% of mCHH sites were 20% methylated. Notably, though the total number of mCG sites was the most similar between C6706 and CV2, their methylation levels were greatly different. The methylation levels at mCG sites changed from 68% mCG sites 10% methylated, 32% mCG sites 20% methylated in C6706 to 19% mCG sites 10% methylated, 57% mCG sites 20% methylated, and 18% mCG sites 30% methylated in CV2 (Fig. 1A).

**Figure 1.**
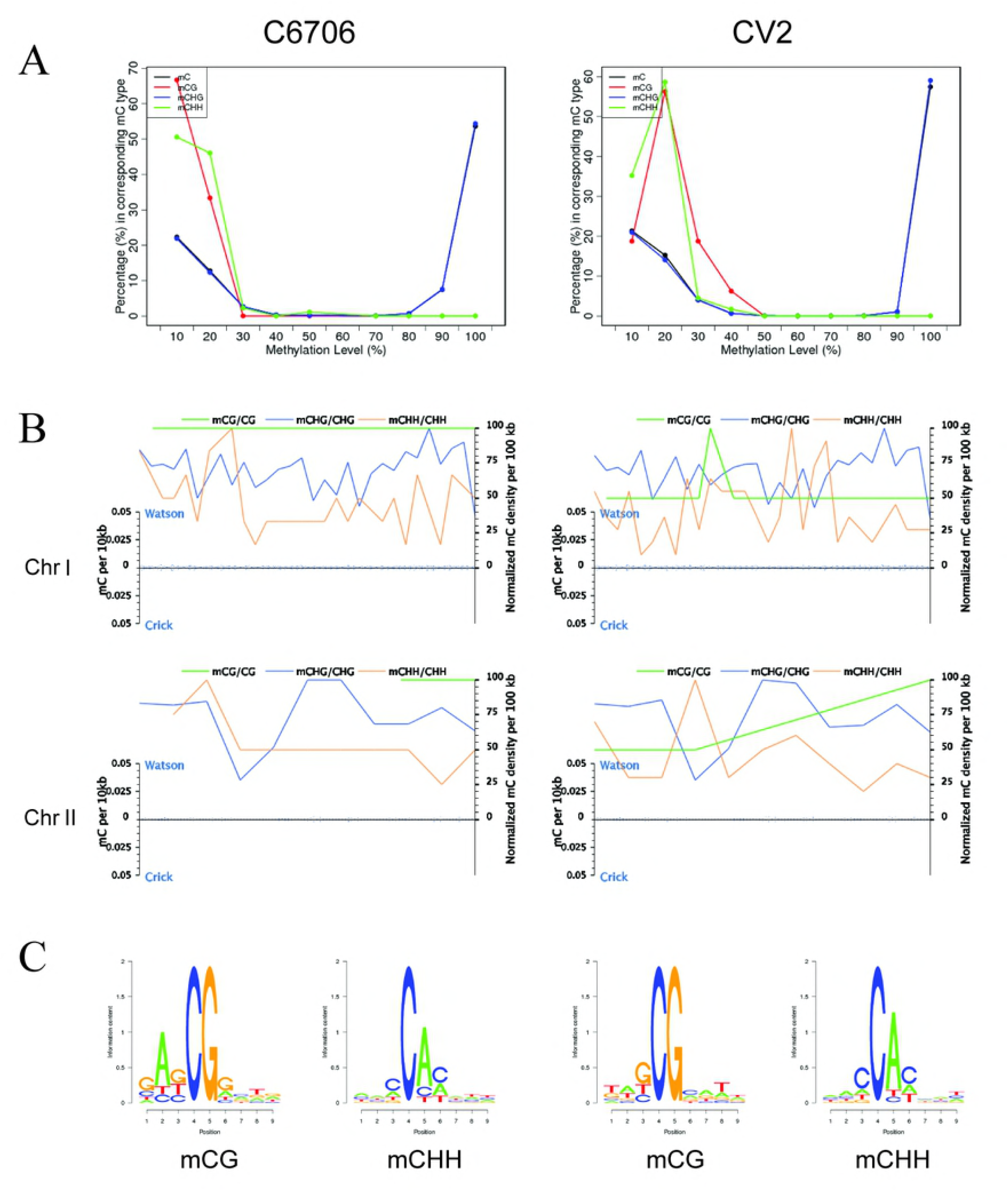
Global trends of the methylomes of C6706 and CV2, and logo plots of the non-CG methylation. A. Distribution of the methylation level in C6706 and CV2 samples. The x axis indicates the methylation level and the y axis indicates the fraction of a specific methylation of mC in all methylcytosines, i.e. the proportion of the sequence in the effective covering sequence on the C base site that supports the site as the methylation site. B. The density distribution of mC on each chromosome. The x-axis represents the chromosome. The y-axis on the left represents mC density calculated from a 10 kb window, and the blue dot represents the distribution of the mC density on the chromosome. The y-axis on the right represents the normalized mC ratio. The curves represent the density distribution of different types of mC bases (CG, CHG and CHH). C. Logo plots of the sequence features of adjacent bases at C site. The x-axis represents the base position, with the C-base for analysis in the fourth position. The y-axis represents the entropy value.

The global-scale view of DNA methylation density in 100 kb windows (Fig. 1B) revealed that there was no correlation between the distributions of three methylation types on the two chromosomes of *V. cholerae*. The density of mCG methylation was relative even within each chromosomal region, while the non-CG methylation showed large variations throughout each chromosome. In C6706 and CV2, the smooth profiles of mCHG density remained constant, but the mCHH density profiles changed dramatically without any similarity (Fig. 1B).

The sequence characteristics of the bases in the vicinity of a methylation site reflects the preference of methylation enzymes to recognition sequences, so we calculated the methylation percentage of the nine bases upstream and downstream of the methylation site (mC at the fourth base). Enrichment of particular local sequences were observed for both mCG and non-CG methylcytosine (Supplementary Fig. 1). 5’-RCCGGY-3’ was the recognition sequence of mCHG and preferences for an AG dinucleotide of mCG and a C of mCHH upstream of the methylation sites were observed. This result was consistent with the analysis of *vchM* [11], but also showed that in addition to the 5’-RCCGGY-3’ sequence, although the 5’-CCWGG-3’ sites were unmethylated, other ^m5^C methylated sequences existed in *V. cholerae*. C6706 and CV2 were identical in this sequence preference, but there were differences in the relative frequency of the bases before and after the recognition sites of mCG and mCHH (Fig. 1C).

In C6706, 6833 ^m5^C methylation sites were located in 1436 genes on chromosome I and in 446 genes on chromosome II, accounting for 49.60% and 39.85% of the total genes of the two chromosomes respectively, and included 459 ^m5^C sites in intergenic regions. In CV2, 6851 ^m5^C methylation sites were located in 1554 genes on chromosome I, 492 genes on chromosome II, and 471 sites in intergenic regions. Between the two samples, 6466 ^m5^C methylation sites kept constant. There were 499 differential ^m5^C sites in C6706, and 856 sites in CV2, involving 377 and 635 genes respectively. There were almost no identical mCHH methylation sites between C6706 and CV2, similar to the mCG sites on chromosome II of the two strains (Supplementary table 2). Overall, DNA methylation in non-CG contexts (mCHG and mCHH) accounted for an absolute majority (99.87% and 99.78%) in *V. cholera* and also accounted for most of the differences in methylation before and after plasmid transformation.

### 2.3 Methylation of host’s MTase genes

Four of the six MTase genes on the *V. cholerae* chromosomes had ^m4^C and ^m6^A methylated sites; VC1769 was the only gene that had all the three methylation types. We did not detect any typed methylation sites on CVA0627 and the orphan Dcm gene, VCA0158 (Table 5). After transformation with pVC211, only a few ^m6^A sites changed, while less than half of ^m4^C sites remained unchanged. Therefore, it was reasonable to observe the large-scale change of methylation profile of the host chromosomes by SMRT sequencing. Combining with changes in ^m5^C sites determined by bisulfite sequencing, we suggested that changes in host methylation profile were partly attributable to the indirect role of host MTases that were affected by the plasmid and partly to the direct role of plasmid MTases themselves.

**Table 5.**
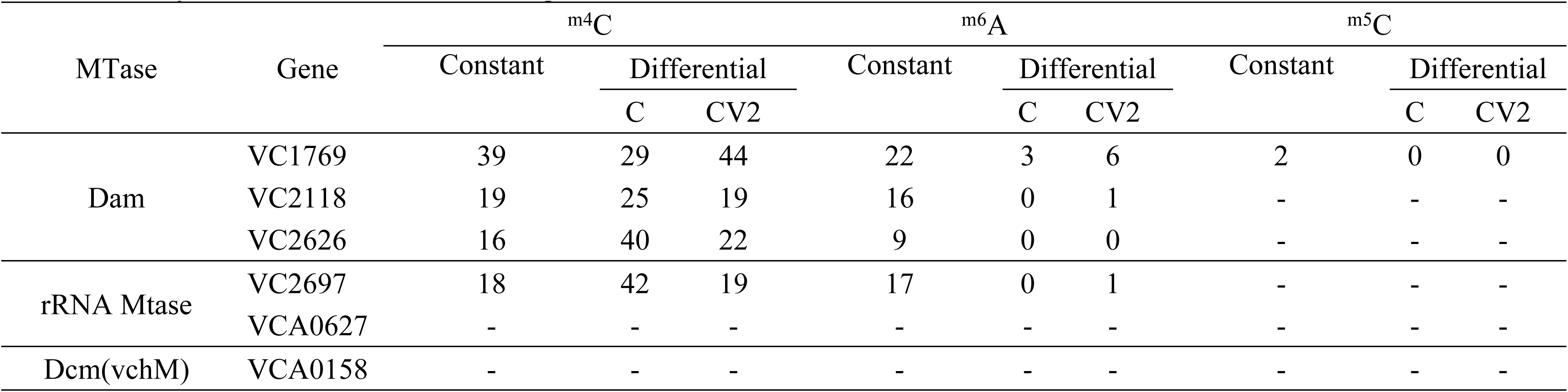
Methylation sites on the MTase genes of the host *V. cholerae* strain.

## 3 Discussion

Methylation plays important roles in epigenetic gene regulation in both eukaryotic and prokaryotic organisms. Compared with the explicit function of Dam methylation in mismatch repair, initiation of chromosome replication, transcription regulation at promoters containing GATC sequences, gene expression, pathogenicity and DNA stable under antibiotic pressure [19], there are limited details known regarding the function of the Dcm MTase [2]. The discovery of two of the three MTase genes on the IncA/C_2_ plasmid were cytosine-specific was intriguing. The original function elucidated for Dcm was to enable the discrimination of self from non-self DNA, as a part of the RM system [20], and to affect plasmid transfer efficiency and dissemination of antibiotic resistance genes in *Enterococcus faecalis* [14]. All of the three types of methylation, ^m6^A, ^m4^C and ^m5^C, and a large amount of methylation without obvious methylation patterns exist in *V. cholerae*. The comparison of genome-wide methylation profiles of *V. cholerae* C6706 before and after the conjugation of pVC211 showed that the MTase genes of the IncA/C plasmid not only affect ^m5^C methylation sites, but also change the number and the characteristics of the recognized sequences of ^m6^A and ^m4^C. The changes involved most of the regions of the two chromosomes, and no obvious preference for gene function was observed. It seemed that the host chromosomes were completely relabeled with a methylation pattern different from that of the host itself. Surprisingly, the new methylation pattern induced by the plasmid was accepted by the restriction enzymes of the host’s RM system. Instead of mimicking the host’s methylation pattern [21], pVC211 actively changed the host’s methylation pattern. In this way, the IncA/C plasmid crossed the barrier of the host’s RM system, demonstrating a novel mechanism by which plasmids of the IncA/C family were able to survival and function in a broad range of host.

## 4 Materials and Methods

### 4.1 Strains and plasmids

*V. cholerae* O1 El Tor strain C6706 was isolated in 1991 from Peru [16]. The IncA/C_2_ plasmid pVC211 was identified in a toxigenic O139 serogroup strain VC211 isolated in 2003, Guangdong province, China [6]. The pVC211 plasmid was transferred into C6706 by conjugation and the strain CV2 was constructed.

### 4.2 Bisulfite sequencing

Genomic DNA was fragmented into ∼250 bp by using Bioruptor (Diagenode, Belgium). The fragments were blunt-ended and phosphorylated, 3’-dA overhung and ligated to methylated sequencing adaptors. Afterwards, samples were bisulfite treated by EZ DNA Methylation-Gold kit (ZYMO), desalted and size selected by 2% agarose gel electrophoresis. After PCR amplification and another round of size selection again, the qualified libraries were sequenced. Sequencing data were filtered and those low-quality data were removed. Then, clean data was mapped onto the reference genome (the complete genome sequences of the standard strain N16961, AE003852 and AE003853; BSMAP) to obtain methylation information of cytosine throughout the whole genome for standard and personalized bioinformatics analysis. The average methylation level was calculated as follows:

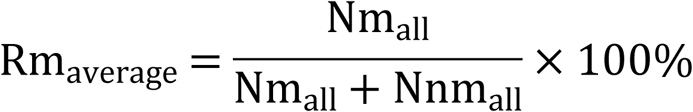

Where Nm is the number of methylated C, and Nnm is the number of reads of non-methylated C. Windows containing at least 5 CG (CHG or CHH) at the same location in the two genomes were identified and the difference in the level of CG methylation in these windows between the two samples were compared to find the regions with significant methylation difference (different methylated region, DMR; 2-fold difference, Fisher test P value ≤ 0.05). If two adjacent DMRs form a contiguous region with significantly different methylation levels in both samples, the two DMRs were merged into one continuous DMR, otherwise they were considered separate DMRs.

CIRCOS was used to compare the differences in the methylation levels of DMRs between samples. The degree of difference of methylation level at one site in two samples was calculated by the following formula, where Rm1 and Rm2 represent the mC methylation level of the two samples, respectively. If the value of Rm1 or Rm2 is 0, it was replaced with 0.001 [17].

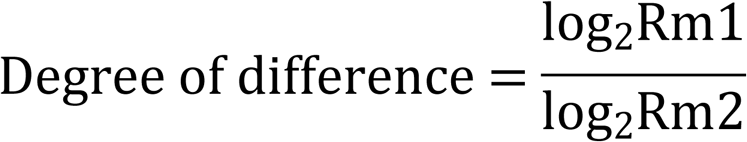

### 4.3 SMRT sequencing

First, the target fragments were amplified from the qualified DNA samples by PCR. Then the damaged ends of the fragments were repaired. Both sides of the DNA fragments were respectively connected with hairpin adapter to get a dumbbell (set of horse ring) structure, which known as SMRTbell. After annealing, the SMRTbell was fixed at the bottom of the ZWM polymerase and used for sequencing. SMRT sequencing was carried out on the PacBioRS instrument (Pacific Biosciences; Menlo Park, CA, USA) using standard protocols. Clean data filtered out polymerase reads with a length less than 50 bp or a mass value less than 0.75 were mapped to the reference genome (AE003852 and AE003853) using the P_Mapping module of SMRT analysis. To identify modified positions, the P_Modification detection module was used. All kinetic raw data in SMRT has been deposited in NCBI (SRA, PRJNA477395).

## Supporting Information Legends

Supplementary Figure 1. The methylation percentage of the nine bases flanking the metnylation site (mC zt the fourth base).

Supplementary Table 1. Genes with obviously changed methylation type in SMRT sequencing

Supplementary Table 2. m5C methylation sites in bisulfite sequencing.

## References

1. Fricke WF, Welch TJ, McDermott PF, Mammel MK, LeClerc JE, et al (2009) Comparative genomics of the IncA/C multidrug resistance plasmid family. J Bacteriol 191: 4750–4757.

2. Harmer CJ, Hall RM (2015) The A to Z of A/C plasmids. Plasmid 80: 63–82.

3. Welch TJ, Fricke WF, McDermott PF, White DG, Rosso ML, et al (2007) Multiple antimicrobial resistance in plague: an emerging public health risk. PLOS One 2: e309.

4. Giske CG, Froding I, Hasan CM, Turlej-Rogacka A, Toleman M, et al (2012) Diverse sequence types of *Klebsiella pneumoniae* contribute to the dissemination of *bla*NDM-1 in India, Sweden, and the United Kingdom. Antimicrob Agents Chemother 56: 2735–2738.

5. Hsueh PR (2010) New Delhi metallo-ss-lactamase-1 (NDM-1): an emerging threat among Enterobacteriaceae. J Formos Med Assoc 109: 685–687.

6. Wang R, Liu H, Zhao X, Li J, Wan K (2018) IncA/C plasmids conferring high azithromycin resistance in Vibrio cholerae. Int J Antimicrob Agents 51: 140–144.

7. Wang R, Yu D, Zhu L, Li J, Yue J, et al (2015) IncA/C plasmids harboured in serious multidrug-resistant *Vibrio cholerae* serogroup O139 strains in China. Int J Antimicrob Agents 45: 249–254.

8. Doublet B, Boyd D, Douard G, Praud K, Cloeckaert A, et al (2012) Complete nucleotide sequence of the multidrug resistance IncA/C plasmid pR55 from *Klebsiella pneumoniae* isolated in 1969. J Antimicrob Chemother 67: 2354–2360.

9. Glenn LM, Englen MD, Lindsey RL, Frank JF, Turpin JE, et al (2012) Analysis of antimicrobial resistance genes detected in multiple-drug-resistant *Escherichia coli* isolates from broiler chicken carcasses. Microb Drug Resist 18: 453–463.

10. Fernandez-Alarcon C, Singer RS, Johnson TJ (2011) Comparative genomics of multidrug resistance-encoding IncA/C plasmids from commensal and pathogenic *Escherichia coli* from multiple animal sources. PLOS One 6: e23415.

11. Banerjee S, Chowdhury R (2006) An orphan DNA (cytosine-5-)-methyltransferase in Vibrio cholerae. Microbiology 152: 1055–1062.

12. Casadesus J, Low D (2006) Epigenetic gene regulation in the bacterial world. Microbiol Mol Biol Rev 70: 830–856.

13. Marinus MG, Lobner-Olesen A (2014) DNA Methylation. EcoSal Plus 6.

14. Huo W, Adams HM, Zhang MQ, Palmer KL (2015) Genome modification in *Enterococcus faecalis* OG1RF assessed by bisulfite aequencing and single-molecule real-time aequencing. J Bacteriol 197: 1939–1951.

15. Murray IA, Clark TA, Morgan RD, Boitano M, Anton BP, et al (2012) The methylomes of six bacteria. Nucleic Acids Res 40: 11450–11462.

16. Thelin KH, Taylor RK (1996) Toxin-coregulated pilus, but not mannose-sensitive hemagglutinin, is required for colonization by *Vibrio cholerae* O1 El Tor biotype and O139 strains. Infect Immun 64: 2853–2856.

17. Heyn H, Li N, Ferreira HJ, Moran S, Pisano DG, et al (2012) Distinct DNA methylomes of newborns and centenarians. Proc Natl Acad Sci U S A 109: 10522– 10527.

18. Egan ES, Waldor MK (2003) Distinct replication requirements for the two *Vibrio cholerae* chromosomes. Cell 114: 521–530.

19. Cohen NR, Ross CA, Jain S, Shapiro RS, Gutierrez A, et al (2016) A role for the bacterial GATC methylome in antibiotic stress survival. Nat Genet 48: 581–586.

20. Takahashi N, Naito Y, Handa N, Kobayashi I (2002) A DNA methyltransferase can protect the genome from postdisturbance attack by a restriction-modification gene complex. J Bacteriol 184: 6100–6108.

21. Bottacini F, Morrissey R, Roberts RJ, James K, van Breen J, et al (2018) Comparative genome and methylome analysis reveals restriction/modification system diversity in the gut commensal Bifidobacterium breve. Nucleic Acids Res 46: 1860– 1877.

